# Estimation of Periodic Signals with Applications to Deep Brain Stimulation

**DOI:** 10.1101/2022.05.23.493124

**Authors:** Paula Chen, Taewoo Kim, Wayne K. Goodman, David A. Borton, Matthew T. Harrison, Jérôme Darbon

## Abstract

Deep brain stimulation (DBS) therapies have shown clinical success in the treatment of a number of neurological illnesses, including obsessive-compulsive disorder, depression, and Parkinson’s disease. An emerging strategy for increasing the efficacy of DBS therapies is to develop closed-loop, adaptive DBS systems that can sense biomarkers associated with particular symptoms and in response, adjust DBS parameters in realtime. The development of such systems requires extensive analysis of the underlying neural signals while DBS is on, so that candidate biomarkers can be identified and the effects of varying the DBS parameters can be better understood. However, DBS creates high amplitude, high frequency stimulation artifacts that prevent the underlying neural signals and thus the biological mechanisms underlying DBS from being analyzed. Additionally, DBS devices often require low sampling rates, which alias the artifact frequency, and rely on wireless data transmission methods that can create signal recordings with missing data of unknown length. Thus, traditional artifact removal methods cannot be applied to this setting. We present a novel periodic artifact removal algorithm for DBS applications that can accurately remove stimulation artifacts in the presence of missing data and in some cases where the stimulation frequency exceeds the Nyquist frequency. The numerical examples suggest that, if implemented on dedicated hardware, this algorithm has the potential to be used in embedded closed-loop DBS therapies to remove DBS stimulation artifacts and hence, to aid in the discovery of candidate biomarkers in real-time. Code for our proposed algorithm is publicly available on Github.

## 1 Introduction

Deep brain stimulation (DBS) is a treatment that regulates neural activity by sending electrical impulses through electrodes implanted in the brain. Currently, DBS is an existing therapy for obsessive compulsive disorder, epilepsy, and Parkinson’s disease, among many other neurological illnesses, but strategies for maximizing the therapeutic effects of DBS while minimizing its side effects remain an active area of research [1]. To that end, one emerging area of interest is the discovery of biomarkers (i.e., neural signals correlated with symptoms) (see, for instance, [2, 3]). The discovery of biomarkers might allow for the development of closed-loop, adaptive DBS systems that are able to sense a biomarker and in response, adjust DBS parameters (e.g., stimulation frequency, amplitude, etc.) in real-time to effectively target symptoms as they arise, while minimizing adverse effects. In order to discover a biomarker and better understand the neurological responses to varying the DBS parameters, the underlying neural signals must be able to be studied while DBS is on. However, DBS creates high amplitude, high frequency stimulation artifacts that contaminate the electrical recordings of the brain. As a result, periodic artifact removal is essential for recovering the underlying neural signal and thus, for understanding the biological mechanisms underlying the qualitative successes of DBS. Furthermore, while biomarker discovery (and thus the corresponding artifact removal) may be done offline, closed-loop, adaptive DBS therapies necessitate the removal of these stimulation artifacts in real-time.

In DBS applications, the artifact removal problem is further complicated by the following: (1) while the stimulation frequency can be set by the DBS device, the device setting provides an estimate of the stimulation frequency that is not accurate enough for successful artifact removal; (2) power constraints of DBS devices often require low sampling rates (e.g., 200-250 Hz), which in turn alias the stimulation frequency near the frequencies of underlying signals of interest; and (3) DBS recordings are often broken into many time segments with unknown phases, e.g., due to modulation of device settings or missing data of unknown length, the latter of which is a common limitation of the wireless data transmission methods used by some DBS devices [4]. While many methods for removing DBS stimulation artifacts exist (see, for instance, [5–9]), no single method, to the authors’ knowledge, addresses all of the aforementioned complications, which are often found in real data. We propose a new method for periodic artifact removal that addresses all of these complications.

The general formulation of our problem of interest is as follows. Given *n* + 1 segments of data, we model the *i*-th segment of the observed signal *S_i_* as:

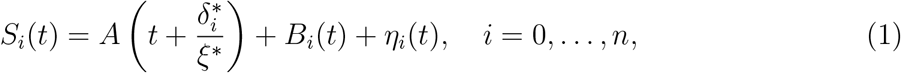

where 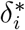 is the (unknown) phase shift between the 0-th and *i*-th segments, *A* is a periodic artifact with (unknown) period 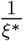, *B_i_* is the recorded neural signal in segment i, and *η_i_* is the noise in segment *i*.

Our goal is to estimate and remove *A* from each *S_i_*. As a result, the underlying signals that we seek to recover are *B_i_* + *η_i_*, *i* = 0,…, n. In practice, *S_i_* is sampled at some collection of discrete times *T_j_*. Then, the following loss function can be used to reconstruct and remove the artifact:

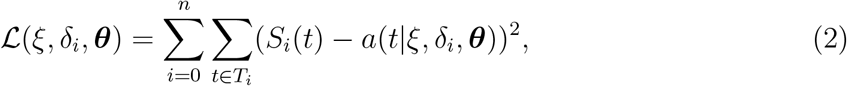

where 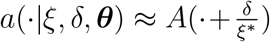 is a model for the artifact that depends on the artifact frequency *ξ*(or equivalently the artifact period 1/*ξ*) and a set (possibly empty) of other parameters ***θ***. Equation (2) generalizes the loss function in [6, 10] to allow for (multiple) unknown phases. In this paper, we consider the following parametric model for the artifact:

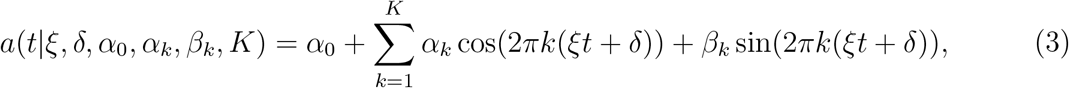

where, in practice, the number of harmonics *K* must be chosen to be finite. Using this parametric model allows for fast computations in our proposed algorithms. Reference [6] uses the same model (3) and [10] experiments with a variety of models for *A*, including (3), but both without phase shifts. Meanwhile, [4] uses the same framework described above to estimate a single unknown phase from two segments (*n* = 1), but does not consider simultaneous inference of the model parameters. Instead, [4] first estimates the artifact and its frequency from a single segment using [6] and then selects among a small finite collection of candidate phases corresponding to an unknown number of missing samples. This process is repeated sequentially to handle multiple phases (*n* > 1). Our method is not sequential; rather, it is based on optimization of (2).

Using the above model (3), a natural way to remove the artifact with the above framework is harmonic regression [11, Chap 2], which for fixed (*ξ*, *δ_i_*) is equivalent to minimizing (2) with respect to ***θ*** = {*α*_0_, *α_k_*, *β_k_*, *k* = 1,…, *K*}. However, in Fig. 1a, we see that harmonic regression requires accurate estimates of the frequency *ξ*. For example, using harmonic regression, 0.001% relative error in the frequency estimate, on average, corresponds to a relative root mean squared error (RMSE) in the reconstructed signal *S_i_* – *a*(·|*ξ, δ_i_, **θ***) of almost 10%. Here, we define relative RMSE as

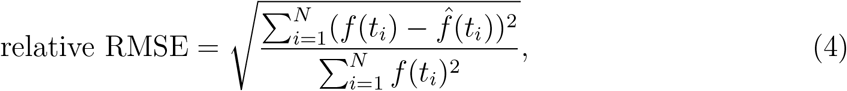

where *f* is the true signal of interest, 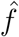 is the estimated signal, and *t_i_* are the sample times. Thus, even somewhat small errors in the frequency can lead to large reconstruction errors.

**Figure 1:**
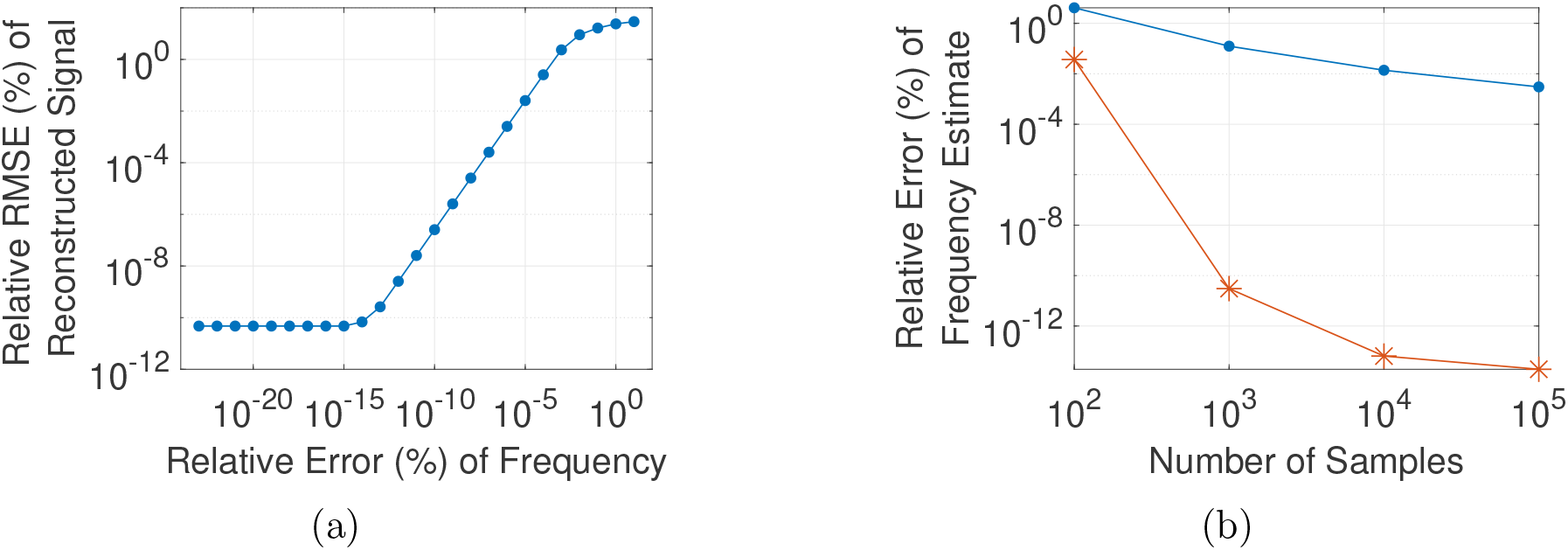
(a) Relative error of frequency vs. relative RMSE of reconstructed signal using harmonic regression. Each point represents the average over 40 trials. For each trial, we use harmonic regression to fit the observed signal with a sinusoid with 5 harmonics and a given frequency estimate. (b) Number of samples vs. relative error of the frequency estimated using a DFT-based method 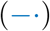 or Algorithm 1 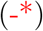. Each point represents the average over 10 trials. In both plots, each trial uses an observed signal sampled at 1000 Hz that consists only of an artifact of the form (3) with no phase shifts and *K* = 5.

Many periodic artifact removal methods rely on the discrete Fourier transform (DFT) to estimate the frequency. However, DFT-based methods generally cannot handle phase shifts and, even segmentwise, are limited in their accuracy for frequency estimation. For example, consider the following simple DFT-based frequency detection method, where the artifact frequency is estimated as the frequency that maximizes the energy (magnitude squared of the DFT) of the observed signal. In Fig. 1b, we observe that even in the case where the observed signal only consists of the periodic artifact and the sampling rate is relatively high (*f_s_* = 1000 Hz), this DFT-based method is unable to accurately estimate the artifact frequency when the frequency does not lie on the DFT grid 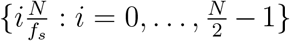, which would be the case for most real-life applications. Specifically, we see that even with 10^5^ samples, this method achieves, on average, greater than 0.001% relative error in the frequency estimate, which, as discussed above, is insufficiently accurate for harmonic regression. In contrast, Fig. 1b shows that our proposed Algorithm 1 achieves much more accurate frequency estimates, e.g., close to machine precision with 10^5^ samples.

### Algorithm 1: Artifact removal algorithm.

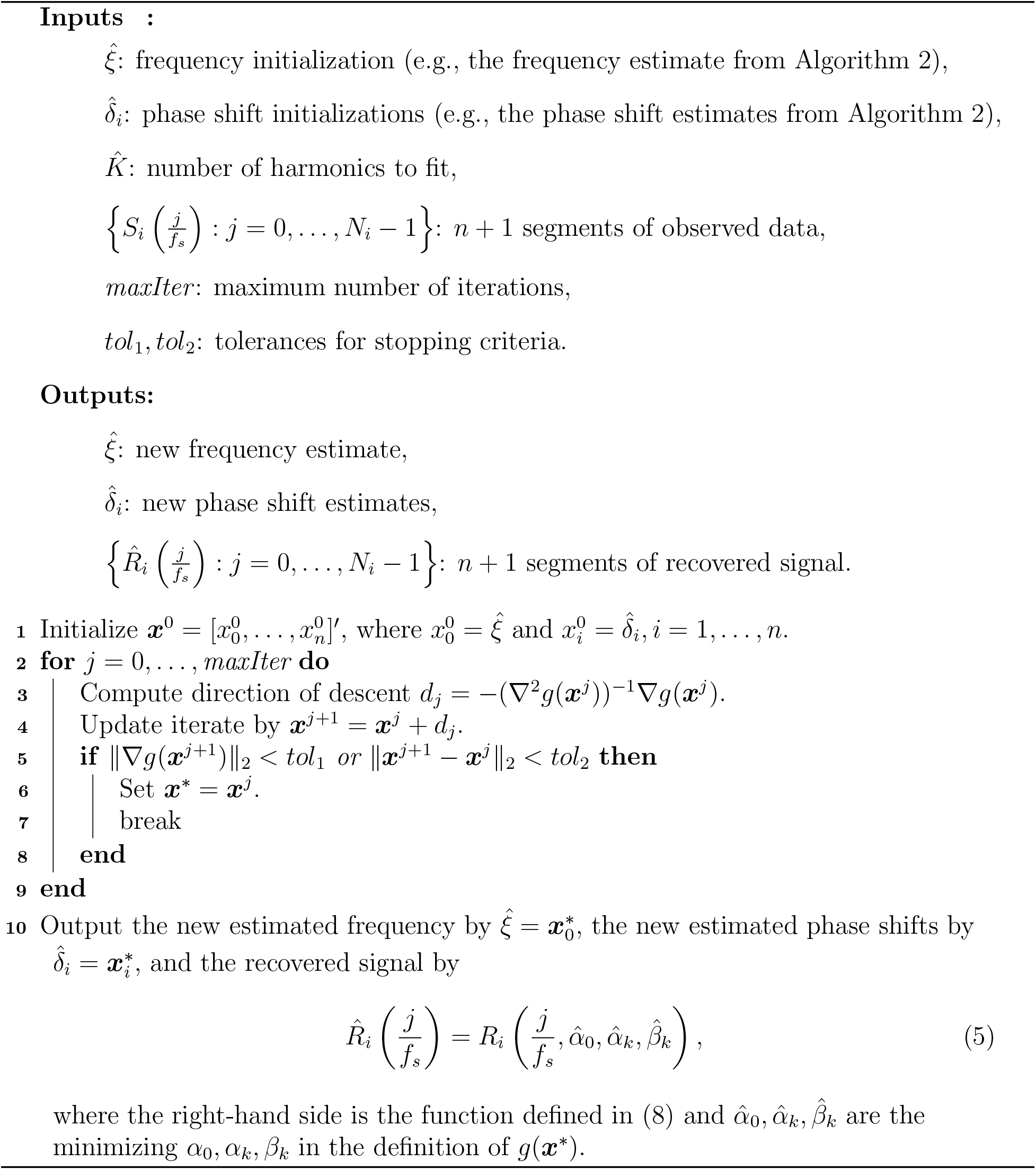

The main contribution of this paper is an iterative periodic artifact removal algorithm that can accurately estimate the artifact frequency in the presence of unknown phase shifts and that involves few tuning parameters. We also provide an initialization algorithm that experimentally provides robust initial estimates of the frequency and phase shifts for our artifact removal algorithm. We demonstrate the performance of our algorithms on the following cases: a simulated artifact with no underlying signal, a simulated artifact with a chirp, an aliased simulated artifact with a simulated underlying signal and missing data, and a human local field potential (LFP) recording. Both algorithms use relatively simple computations and demonstrate relatively short runtimes (in comparison to the length in time of the data processed), which show their potential to be used in embedded closed-loop DBS therapies in real-time.

The remainder of this paper is organized as follows. In Section 2, we describe the details of our method. In Section 2.1, we describe our artifact removal method, which fits the observed signal using harmonic regression while jointly estimating the frequency and phase shifts via a least squares regression. In Section 2.2, we provide an initialization algorithm for the method described in Section 2.1. In Section 3, we provide some numerical examples, which demonstrate some of the capabilities of our proposed method. In Section 4, we provide some concluding remarks. Finally, in Appendix A–B, we provide some explicit formulas for our algorithms.

## 2 Proposed Algorithm

In this section, we describe our algorithm. First, we describe our artifact removal algorithm (Algorithm 1), which removes the artifact by estimating the frequency and phase shifts while jointly fitting the artifact using harmonic regression. Second, we describe an initialization algorithm (Algorithm 2), which finds an initial estimate of the frequency and phase shifts via an energy maximization method. The code for both algorithms and implementation details are available at https://github.com/pxchen95/artifact_removal.

### Algorithm 2: Initialization algorithm for Algorithm 1.

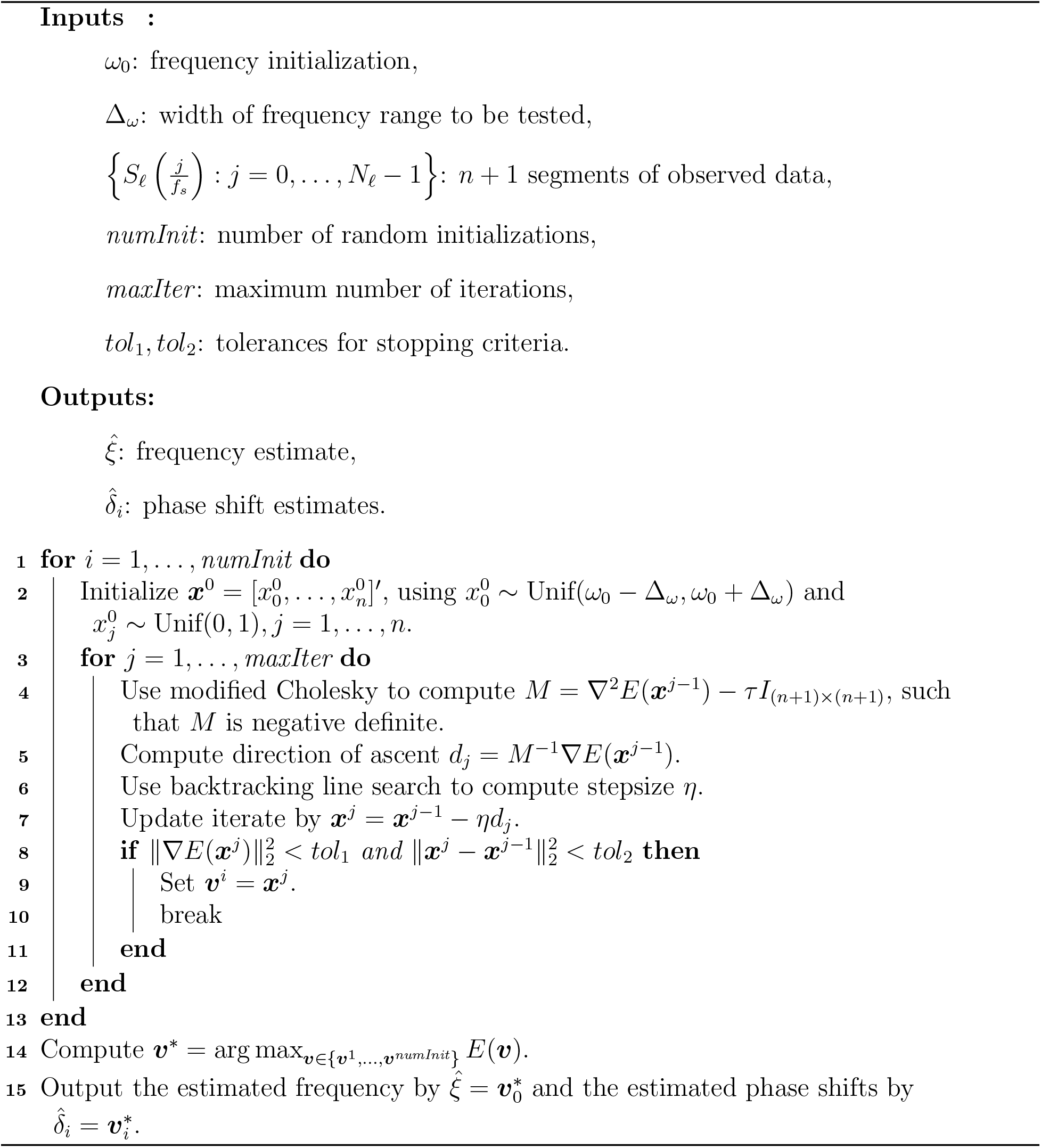

Our model is as follows. We observe *n* +1 segments of data *S_i_, i* = 0,…, *n*, where each *S_i_* is defined by (1) and 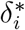 is the (unknown) true phase shift between the 0-th and *i*-th segments (by convention, 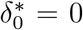). We assume that the artifact can be represented by (3), where the true fundamental frequency *ξ*\*, the true amplitudes 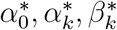, and the true number of harmonics *K* are unknown. A key assumption for our method is that the energy of the artifact at its fundamental frequency and harmonics is sufficiently larger than the energy of the brain at those frequencies (or at the frequencies that they are aliased to). In practice, we do not have access to the continuous signals and only access each *S_i_* at some discrete sample times. While these sample times may be irregularly spaced, we only consider uniformly spaced sample times as, in practice, most DBS devices use uniformly spaced sampling based on relatively stable system clocks; i.e., we assume that the i-th segment is sampled at the times 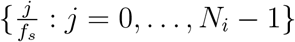, where *f_s_* is the sampling rate.

### 2.1 Artifact Removal Algorithm

Based on the assumed form of the artifact (3), harmonic regression [11, Chap 2] is a natural choice for removing the artifact given an estimate of the frequency. However, as shown in Figure 1a, small perturbations in the frequency estimate can result in large errors in the signal reconstructed by harmonic regression. To mitigate this issue, our method fits the observed signal using harmonic regression while jointly estimating the frequency and phase shifts. Specifically, we remove the artifact by solving the following optimization problem:

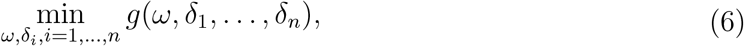

where the objective function *g* is defined by:

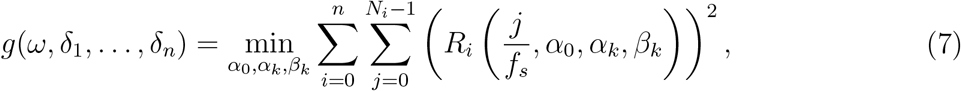

and we denote the i-th segment of the recovered signal by:

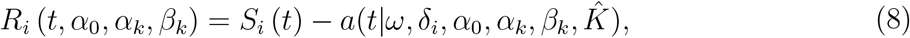

where 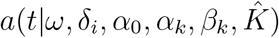 is defined by (3). For each fixed (*ω*, *δ*_1_,…, *δ_n_*), *g*(*ω*, *δ*_1_,…,*δ*_n_) is the result of using harmonic regression to remove a periodic artifact of the form (3) with 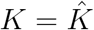 harmonics, fundamental frequency *ξ*\* = *ω*, and phase shifts *δ_i_* from the 0-th observed segment *S*_0_. For our algorithms, the number of harmonics to fit 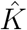 is effectively the only parameter that needs to be tuned. We note that minimizing over *α*_0_ in (7) means that the recovered signal will have mean 0. Including this amplitude is optional, and if *α*_0_ is omitted in (8), then the recovered signal may have nonzero mean, but the reconstructed artifact will necessarily have mean 0. We also note that for each fixed (*ω, δ*_1_,…, *δ_n_*), *g*(*ω, δ*_1_,…,*δ_n_*) can then be computed exactly up to numerical precision errors. Specifically, we set the gradient of the objective function in (7) to zero and solve the resulting linear system using any appropriate numerical linear algebra solver.

In Algorithm 1, we use a Newton’s descent method to solve (6). The numerical complexity of Algorithm 1 is dominated by the computation of the descent direction, which requires us to solve an (*n* +1) × (*n* +1) and a 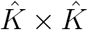 linear system. Thus, the numerical complexity of the algorithm is 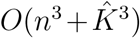. In practice, we expect both *n* and 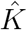 to be relatively small since the data can always be segmented so as not to consider too many gaps at a time and many DBS devices have built-in low-pass filters that effectively bandlimit the recorded signals. Note that we can apply Newton’s descent to this problem since *g* is twice differentiable, where the differentiability of *g* follows from [12, Chap 4.3, Thm 4.13]. Also note that while *g* is not convex, it is locally convex at the global minimizer. Hence, the initialization for Algorithm 1 is crucial for convergence. In the next section, we introduce an initialization algorithm (Algorithm 2) for Algorithm 1. Based on our numerical experiments, we found that Algorithm 2 typically has relatively short computational runtime and provides sufficiently accurate initializations for Algorithm 1.

### 2.2 An Initialization Algorithm

In this section, we present an initialization algorithm for Algorithm 1. Specifically, our algorithm locally maximizes the energy of the observed signal using discrete samples but without being constrained to frequencies on the DFT grid.

Define the Fourier transform for our problem as

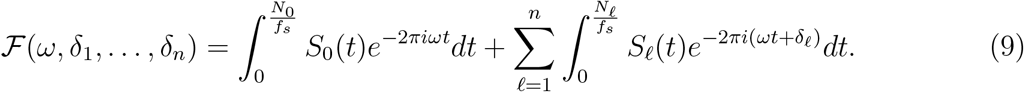

We interpret (9) as trying to align all of the segments *S_ℓ_* (*ℓ* = 0, 1,…, *n*) in time to be in phase with a single wave *e*^−2*πiωt*^. Next, define the energy as

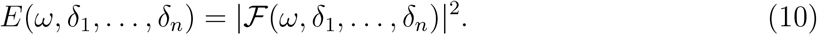

By straightforward calculation, it is easy to verify that *E* is twice differentiable and nonconcave. We propose to recover the true frequency *ξ*\* and true phase shifts 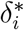 by solving the following optimization problem:

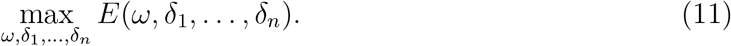

Note that for fixed *ω, δ_j_*, *j* ≠ *i*, the function 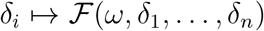 (and hence, the function *δ_i_* ↦ *E*(*ω, δ*_1_,…,*δ_n_*)) is 1-periodic. Thus, solving (11) can only recover the phase shifts 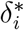 up to a multiple of 1. Moreover, since E is nonconcave, the maximizers of (11) are not unique.

In Algorithm 2, we solve (11) numerically by applying a modified Newton’s ascent method (based on modified Cholesky [13]) with backtracking line search and random initialization. Since we only have access to discrete samples (and not the continuous signals), we approximate all of the integrals in the definitions of the energy *E* (10), its gradient Δ*E*, and its Hessian Δ^2^*E* using trapezoidal rule [14, Chap 8.1]. The computational effort for each iteration of Algorithm 2 is proportional to (*n* + 1)^3^. As before, we note that, in practice, the data can always be segmented so that n is relatively small. Additionally, the implementation of Algorithm 2 can be made more efficient by parallelizing the computations for each random initialization.

## 3 Results

In this section, we present several numerical examples that demonstrate some of the capabilities of our proposed algorithms. In examples 1-3, we set the artifact *A* to be of the form (3) with 5 harmonics and true fundamental frequency *ξ*\* ≈ 150.6117 Hz. In example 4, we use a human local field potential (LFP) dataset, where the stimulation frequency is set by the DBS device to 150.6 Hz. Examples 3-4 contain missing data. For these examples, the phase shifts *δ_i_* correspond to the time gaps in the data, and we define the segments *S_i_* to be the contiguous segments of observed data. In all of the examples, we remove the reconstructed artifact from *S_i_* to recover the underlying signals *B_i_* + *η_i_* using Algorithm 1 with 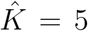, *maxiter* = 1000, *tol*_1_ = 1*e* – 8, and *tol*_2_ = 1*e* – 16 and initialized using the outputs of Algorithm 2 with *ω*_0_ = 150.6, Δ*_ω_* = 5, *numInit* = 25, *maxiter* = 5000, and *tol*_1_ = *tol*_2_ = 1*e* – 8. Since Examples 1-3 all use simulated data, we report the corresponding results in arbitrary units (a.u.); we note that, unless stated otherwise, the scaling of the units are consistent between the subfigures of each figure corresponding to these simulated examples. We run all of the numerical examples using a simple MATLAB implementation on an 8th Gen Intel Laptop Core i5-8250U with a 1.60GHz processor.

### 3.1 Example 1: A Simulated Artifact with No Underlying Signal and No Phase Shifts

In this example, we set the underlying signals *B*_0_ = *η*_0_ = 0 and use the sampling rate *f_s_* = 1000 Hz, 10^4^ samples, and no phase shifts (*n* = 0). Since there are no phase shifts in this example, it is possible to initialize Algorithm 1 using the output of other existing frequency estimation algorithms. However, we use the initialization from Algorithm 2 to demonstrate our algorithms’ ability to perform basic frequency detection and to visualize the key functions being optimized by our algorithms.

Figure 2a depicts plots of the energy *E*, which is computed using trapezoidal rule to compute the integrals in (10). In Figure 2a, we see that *E* is nonconcave and that Algorithm 2 converges to the numerical maximizer of *E*, but the approximation errors resulting from the trapezoidal rule cause the numerical maximizer of *E* to be slightly offset from the true frequency of the artifact. Figure 2b depicts plots of the least squares objective function *g*, as defined by (7). In Figure 2b, we see that *g* is nonconvex and that Algorithm 1 converges to the true frequency and obtains a much more accurate estimate in comparison to Algorithm 2. Specifically, the frequency estimate resulting from Algorithm 1 has a relative error of 3.7742 × 10^-14^%, whereas the frequency estimate resulting from Algorithm 2 has a relative error of 3.2337 × 10^-6^%. In Figure 2c, we observe that the artifact reconstructed by our algorithms matches the true artifact very well with a relative RMSE of 1.7918 × 10^-10^%. In Figure 2d, we see that the signal recovered by our algorithms is approximately 0 with an RMSE of 3.0106 × 10^-10^. Finally, in Figures 2e and 2f, we observe that in the frequency domain, the artifact is no longer detectable in the recovered signal.

**Figure 2:**
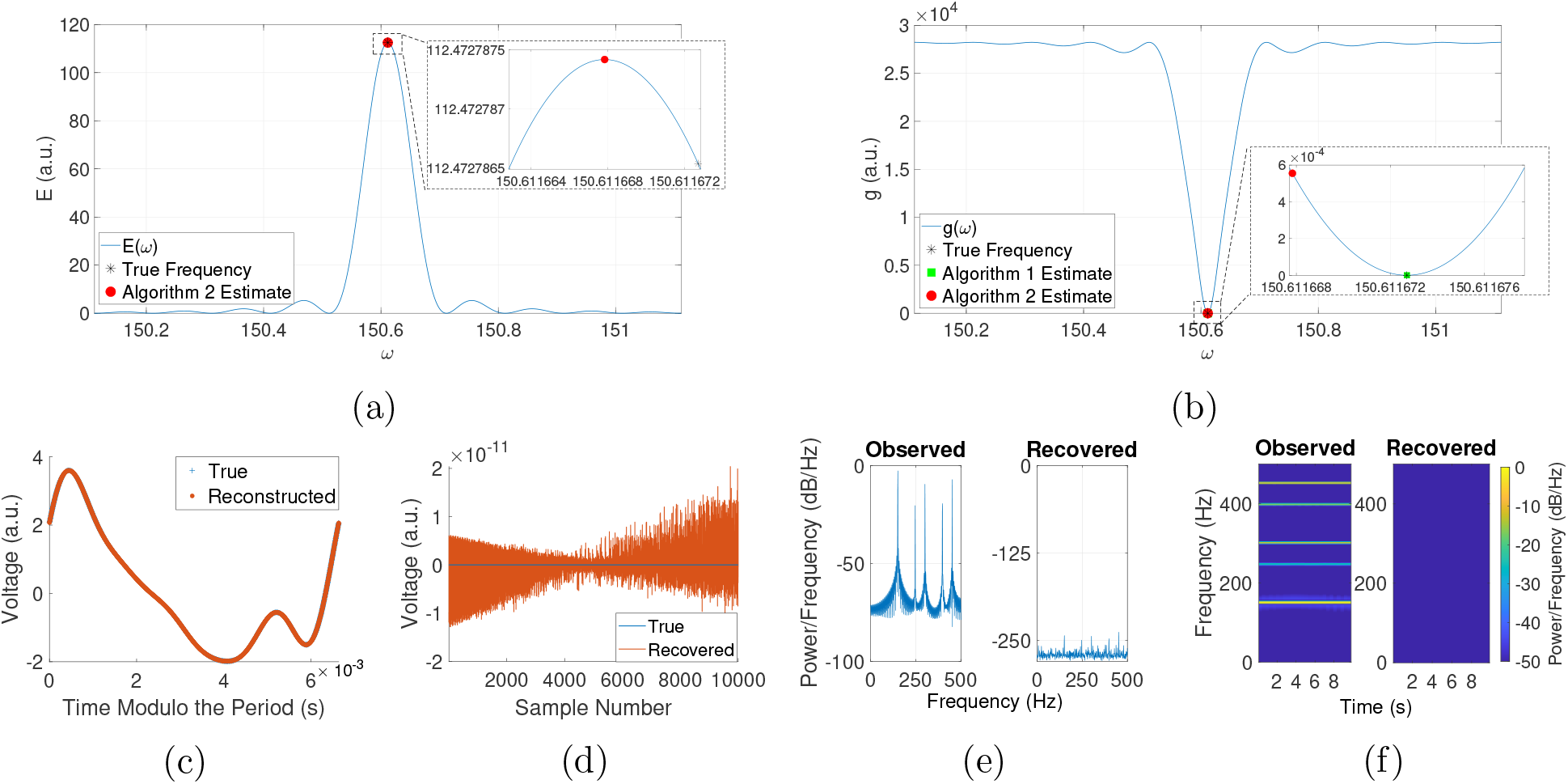
Removing a well-sampled artifact with no underlying signal. (a) Plot of the energy *E*, as computed using trapezoidal rule. (b) Plot of the least squares objective function *g*, as defined by (7). (c) Plot of the true and reconstructed artifacts (with time modulo the true and Algorithm 1 estimate of the period, respectively). (d) Plot of the true and recovered underlying signals. (e) Welch’s power spectral density. (f) Spectrogram using short-time Fourier transforms. (a.u. = arbitrary units)

### 3.2 Example 2: A Simulated Artifact with a Chirp and No Phase Shifts

In this example, we set the underlying signals *B*_0_ to be a chirp with frequencies ranging from 0 Hz to 500 Hz and *η*_0_ = 0 and use the sampling rate *f_s_* = 1000 Hz, 10^4^ samples, and no phase shifts (*n* = 0). We use this example to demonstrate the performance of our algorithms in the presence of underlying signals consisting of short snippets with frequencies at or near the frequency of the artifact.

In Figure 3a, our reconstructed artifact matches the true artifact with a relative RMSE of 0.5837%, where our frequency estimate has relative error 7.7068 × 10^-6^%. In Figure 3b, the recovered underlying signal matches the true underlying signal with a relative RMSE of 5.5508%. In Figure 3c, we see that most of the power at the artifact frequencies is removed by our algorithms, but some small peaks are present near the frequencies corresponding to the artifact harmonics. Similarly, in Figure 3d, we observe that the artifact is mostly removed (the difference in the power at the artifact frequencies in the observed signal versus in the recovered signal is about 50 dB), while the power of the chirp remains intact. These results show that our algorithms are able to accurately recover the underlying signal even when it contains frequencies at or near the artifact frequency.

**Figure 3:**
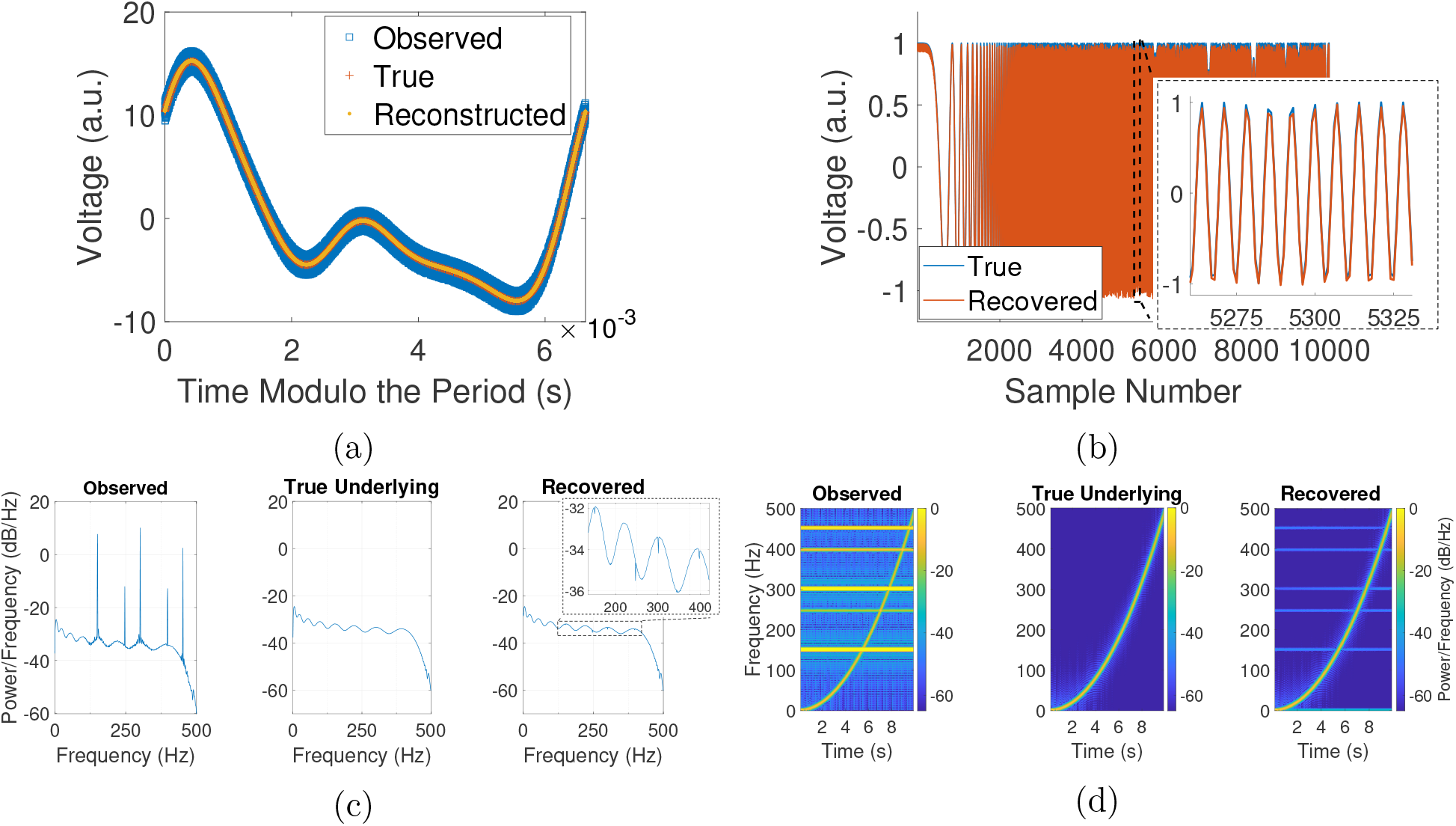
Removing a well-sampled artifact from a chirp. (a) Plot of the observed signal and reconstructed artifact (with time modulo the period estimated by Algorithm 1) and the true artifact (with time modulo the true period). (b) Plot of the true and recovered underlying signals. (c) Welch’s power spectral density. (d) Spectrogram using short-time Fourier transforms. (a.u. = arbitrary units)

### 3.3 Example 3: An Aliased Simulated Artifact with a Simulated Neural Signal, Noise, and Missing Data

In this example, we set the underlying signals *B_i_* to be a simulated neural signal (computed as the sum of short snippets of sinusoidal waves with random frequencies and random lengths and demeaned) and *η_i_* to be independent and identically distributed (iid) Gaussian noise and use the sampling rate *f_s_* = 250 Hz. To simulate missing data, we create 10 segments of contiguous data containing 250 samples each, where the gaps between segments are of random length. Recall that many real-life applications require low sampling rates to handle the battery constraints of DBS devices. In these cases, the fundamental frequency of the stimulation artifact often exceeds the Nyquist frequency [14] and is aliased to potential frequency ranges of interest. Here, the fundamental frequency of the artifact is aliased to approximately 99.3883 Hz. Additionally, recall that the recording methods used by DBS devices can lead to missing data, where the length of the missing data is unknown and the length of the segments of contiguous data is relatively short. Thus, we test the performance of our algorithms in this worst-case scenario, which includes aliasing, short segments of recorded data, and missing data.

In Figure 4a, the reconstructed artifact matches the true artifact with a relative RMSE of 5.5521%, while our frequency estimate has relative error 2.3023 × 10^-3^%. In Figure 4b, we see that in the time domain, the recovered underlying signal matches the true underlying signal with a relative RMSE of 11.0553%, but the two signals visually match very well despite the relative RMSE being somewhat high. Note that in Figure 4b, we overlay each sample of the recovered signal on top of the corresponding sample of the true underlying signal. In other words, the length of the gaps in the recovered signal in this figure do not necessarily correspond to the lengths estimated by Algorithm 1 (recall that Algorithm 1 can only recover phase shifts up to some multiple of 1). In Figure 4c, we observe that in the frequency domain, the power of the artifact frequencies is no longer detectable in the recovered signal, and the recovered signal and true underlying signal are visually indistinguishable. These results show that our algorithms are able to remove stimulation artifacts even in this worst-case scenario.

**Figure 4:**
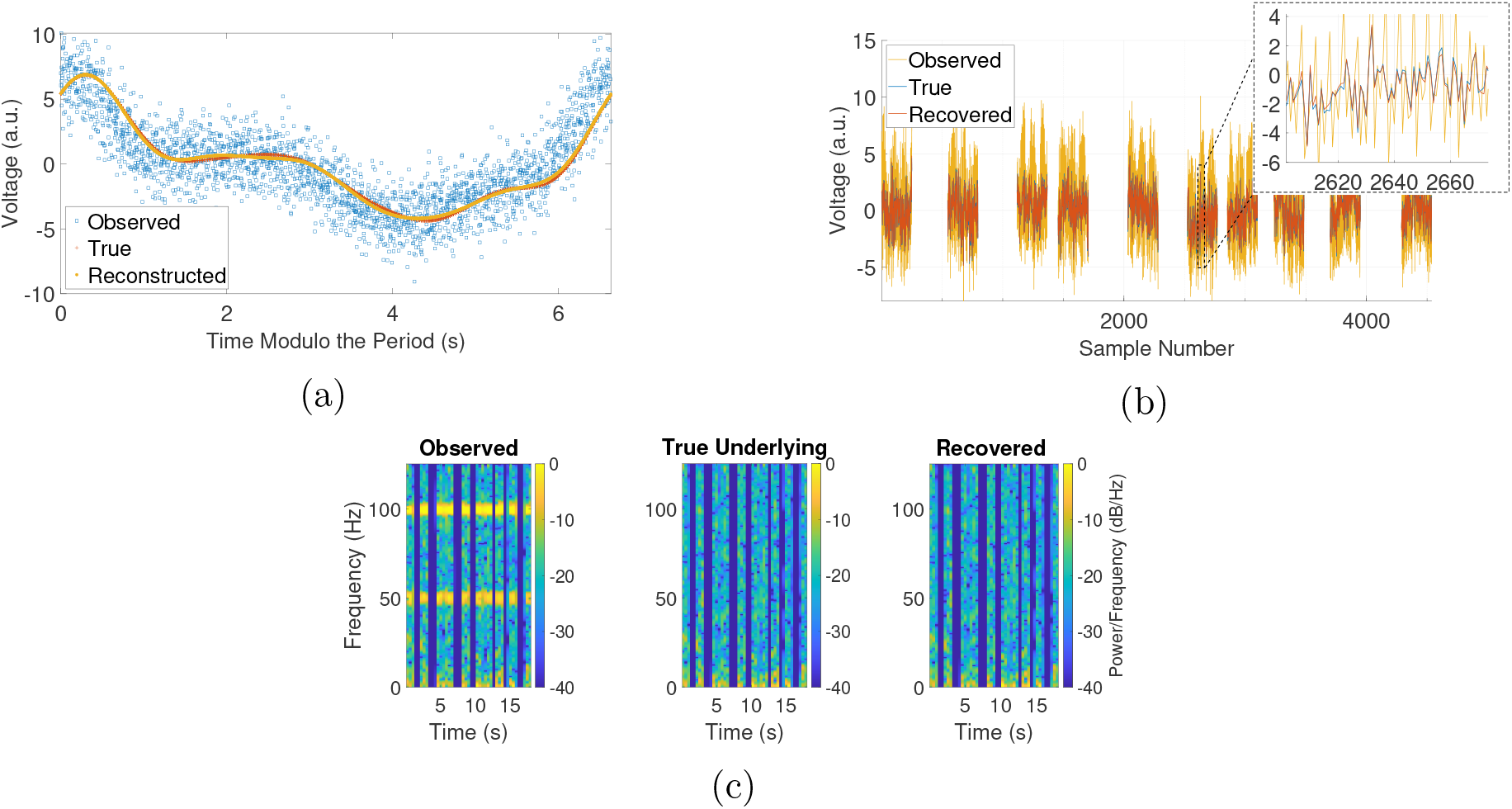
Removing an aliased artifact from a simulated neural signal with missing data. (a) Plot of the observed signal and reconstructed artifact (with time modulo the period estimated by Algorithm 1) and the true artifact (with time modulo the true period). (b) Plot of the true and recovered underlying signals. (c) Spectrogram using short-time Fourier transforms. (a.u. = arbitrary units)

### 3.4 Example 4: Human LFP Recording with Missing Data

In this example, we consider a real recording of a human LFP signal with DBS from a clinical trial (ClinicalTrials.gov #NCT04281134). The research participant (37y/o female) had a history of long-standing OCD and underwent clinically indicated DBS surgery for treatment of OCD using a Summit RC+S (Medtronic, Minneapolis, MN, USA) device. DBS leads (Model 3778) were intracranially placed bilaterally in the VC/VS. The participant gave fully informed consent according to study sponsor guidelines, and all procedures were approved by the local institutional review board at Baylor College of Medicine (H-40255, H-44941, 5/4/2021). LFP was sensed with bipolar contacts around the stimulation contact at a sampling rate of *f_s_* = 1000 Hz. The stimulation frequency was set by the device to 150.6 Hz. We simulated missing data by segmenting the recorded data into 10 segments of contiguous data containing 250 samples each with gaps between the segments that are of random length.

In this example, Algorithm 1 gives us a frequency estimate of approximately 150.6093. In Figure 5a, we see that using our estimates of the artifact period and the time gaps, the observed signal generally looks like a noisier version of our reconstructed artifact. In Figure 5b, we observe that the bands corresponding to the frequencies of the DBS artifact are removed by our algorithm. Note that in this figure, the “No DBS” plot cannot be directly compared with the “Recovered” plot since they each represent signals that were recorded at different times and under different conditions (i.e., with DBS off or on, respectively); however, since we do not have access to the true underlying signal, the former plot provides a baseline for what the plot for the true artifact-free underlying signal may look like. These results demonstrate that our algorithms work on real datasets of interest, even in the presence of many gaps of unknown length.

**Figure 5:**
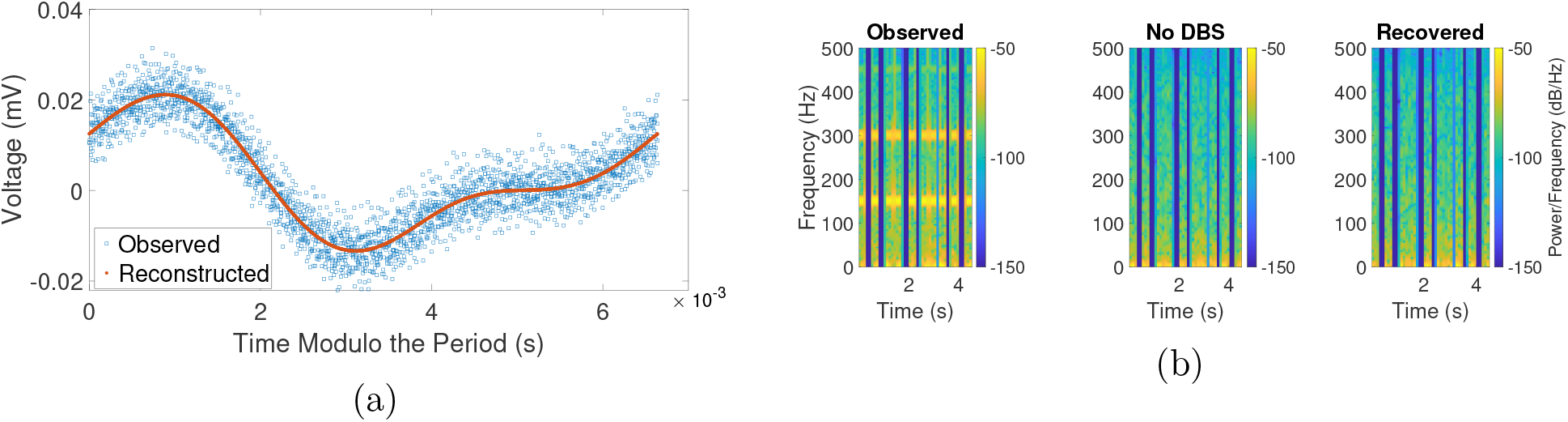
Removing a real DBS artifact from a human LFP recording with missing data. (a) Plot of the observed signal and reconstructed artifact (with time modulo the period estimated by Algorithm 1). (b) Spectrogram using short-time Fourier transforms. Note that the “Recovered” and the “No DBS” plots are not directly comparable as they represent signals that were recorded at different times and under different conditions (i.e., with DBS on or off, respectively).

## 4 Summary

We have proposed a periodic artifact removal algorithm (and a corresponding initialization algorithm) that is able to accurately remove stimulation artifacts in the presence of unknown phase shifts (e.g., missing data of unknown length) and in some cases where the stimulation frequency exceeds the Nyquist frequency. Our algorithms effectively require only one tuning parameter – the number of harmonics 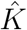 to fit. Choosing 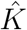 too small corresponds to underfitting the artifact, while choosing 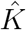 too large corresponds to overfitting the artifact. Thus, the parameter 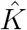 is easily interpreted and could be roughly estimated by counting the number of bands corresponding to the harmonics of the artifact that are visible in a spectogram of the observed signal. However, we also observed that picking a slightly higher 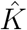 makes little difference in the results. For instance, if we instead use 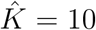 in Example 1 (Section 3.1), then all of the resulting errors are identical to the corresponding reported errors up to three decimal places; meanwhile, using 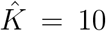 in Example 3 (Section 3.3) results in relative errors that differ from the corresponding reported values by less than 1%. Additionally, while the numerical examples provided in this paper provide promising results, they are not exhaustive. For example, many real-life DBS artifacts consist of semi-rectangular waves (i.e., have infinitely many harmonics), which were not considered in this paper and could be explored in a future study.

Since our algorithms are relatively simple computationally, they have the potential for efficient implementation on dedicated hardware, such as digital signal processors (DSPs) or field-programmable gate array (FPGAs). In each of the examples in Section 3, each of which corresponds to roughly 4-18 seconds of DBS recording, our simple MATLAB implementation of Algorithm 1 took 0.6-1.5 seconds to run, while Algorithm 2 took 0.4-1.0 seconds to run. However, these runtimes would be greatly shortened by more efficient implementations of our algorithms. As such, our algorithms have the potential to be used in embedded closed-loop DBS therapies to remove stimulation artifacts and thus to discover candidate biomarkers in real-time. Our algorithms may also be applicable to more general settings. For example, we defined our algorithms using uniform sample times, but they could be easily extended to handle nonuniform sample times. Furthermore, (7) could be reformulated to minimize over the mean *α*_0_ segmentwise to handle cases where the mean of the underlying signal varies between segments.

## A Some Explicit Formulas for Algorithm 1

In this section, we provide some explicit formulas for various quantities that are needed to implement Algorithm 1. For each fixed (*ω, δ*_1_,…, *δ_n_*), the minimizing *α*_0_, *α_k_*, *β_k_* in *g*(*ω, δ*_1_,…, *δ_n_*) (as defined by (7)) can be computed as the solution to the following lin-ear system:

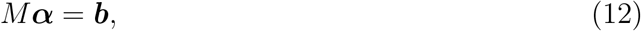

where *M* is a 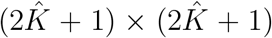 matrix *M*, where the (*i*, *j*)-th element *m_ij_* (for 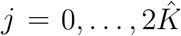) is defined by:

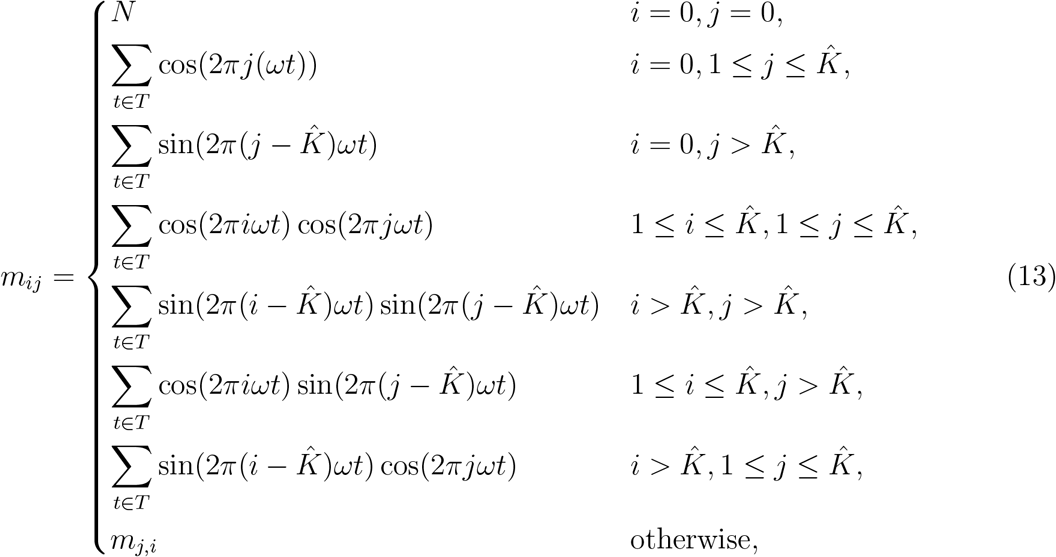

where 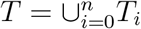 and 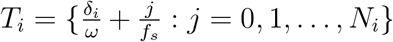, ***b*** is a 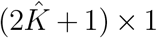 vector, where the *i*-th element *b_i_* (for 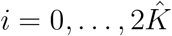) is defined by:

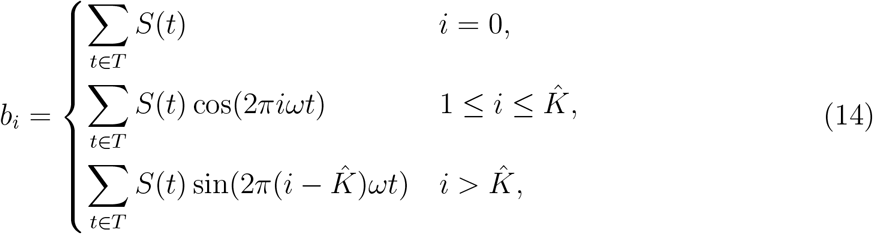

and ***α*** is a 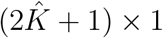 vector, where the *i*-th element ***α_i_*** (for 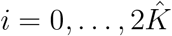) is given by:

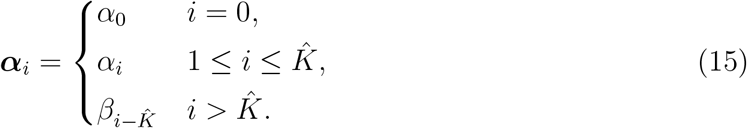

Let 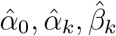 be the minimizing *α*_0_, *α_k_*, *β_k_* in the definition of *g*(*ω*, *δ*_1_,…, *δ_n_*). In what follows, for simplicity of notation, we drop the dependence of the following functions on (*ω*, *δ*_1_,…, *δ_n_*). The first derivatives of g are given by:

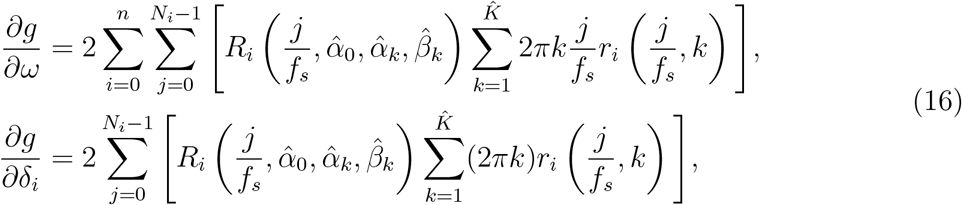

where *i* = 1,…, *n*, *R_i_* is the function defined by (8), and *r_i_* is defined as follows:

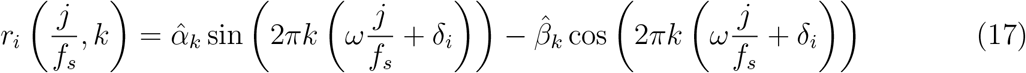

The second derivatives of *g* are given by:

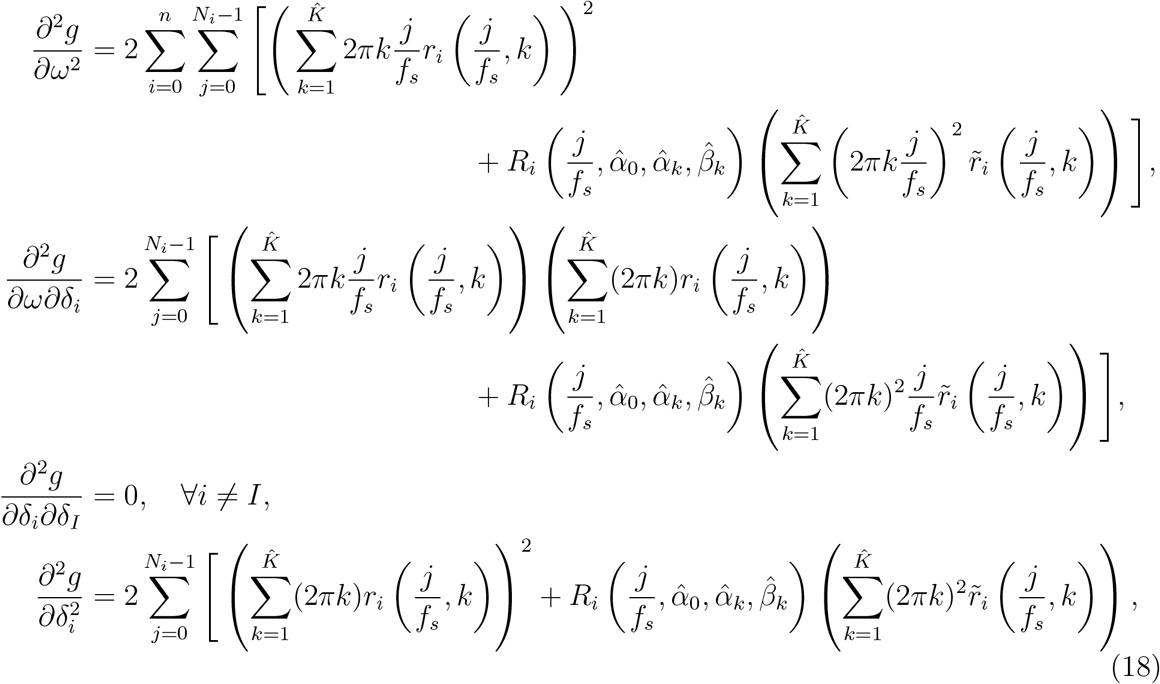

where *i*, *I* = 1,…, *n, R_i_* is the function defined by (8), *r_i_* is the function defined by (17), and 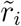 is defined as follows:

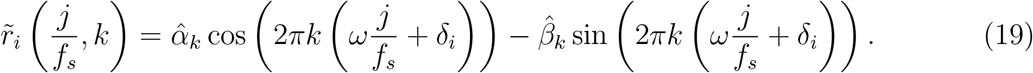

## B Some Explicit Formulas for Algorithm 2

In this section, we provide some explicit formulas for various quantities that are needed to implement Algorithm 2. In what follows, for simplicity of notation, we drop the dependence of the following functions on (*ω, δ*_1_,…, *δ_n_*).

Define 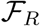 and 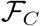 to be the real and imaginary parts of 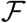 (as defined by (9)), respectively, which can be approximated using trapezoidal rule as follows:

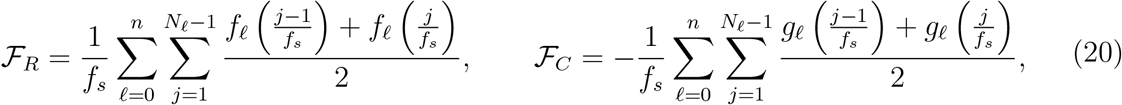

where

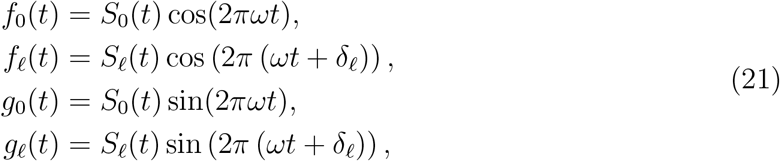

for *ℓ* = 1,…, *n*. Then, we can equivalently compute the energy (originally defined in (10)) as

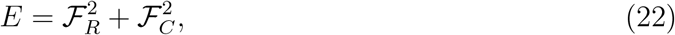

where 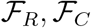 are defined in (20) above. We compute the *i*-th component of ∇*E* as follows:

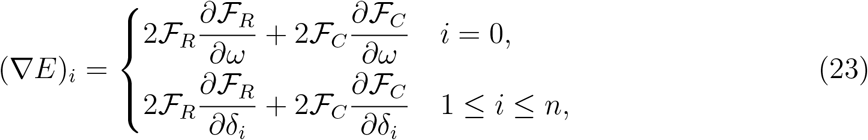

where we compute 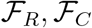 using (20) and we approximate the first-order derivatives of 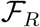 and 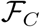 using trapezoidal rule as follows:

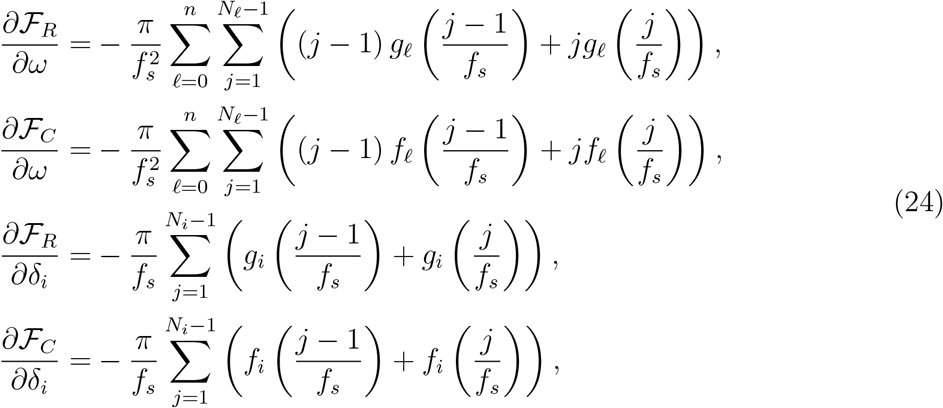

where *f_i_*, *g_i_* are the functions defined in (21). We compute the (*i*, *j*)-th element of ∇^2^*E* as follows:

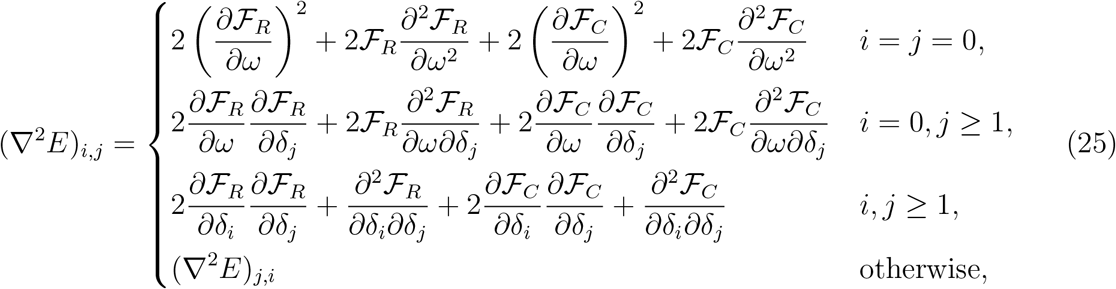

where 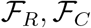 are computed using (20), the first-order derivatives of 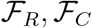 are computed using (24), and the second-order derivatives of 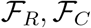 are approximated using trapezoidal rule as follows:

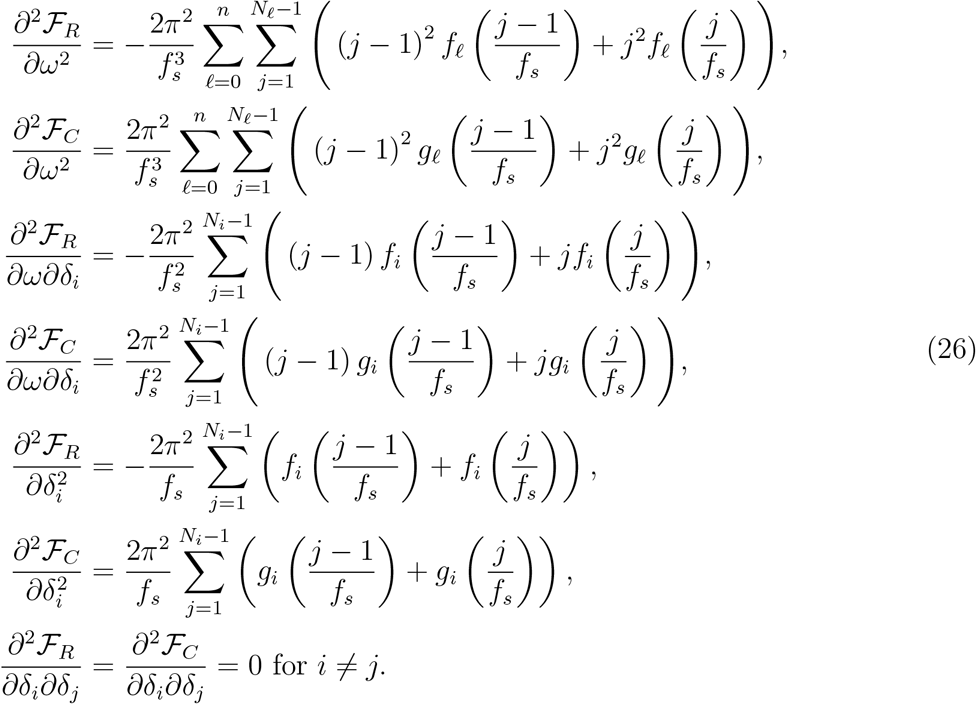

## Acknowledgments

We thank the Open Mind Consortium (https://openmind-consortium.github.io/) for their expertise and resources.

